# The frailty index is a predictor of cause-specific mortality independent of familial effects from midlife onwards

**DOI:** 10.1101/486845

**Authors:** Xia Li, Alexander Ploner, Ida K Karlsson, Xingrong Liu, Patrik KE Magnusson, Nancy L Pedersen, Sara Hägg, Juulia Jylhävä

## Abstract

**Background:** Frailty index (FI) is a well-established predictor of all-cause mortality, but less is known for cause-specific mortality and whether familial effects influence the associations. Furthermore, the population mortality impact of frailty remains understudied.

**Objectives:** To estimate the predictive value of frailty for all-cause and cause-specific mortality, and to test whether the associations are time-dependent. We also assessed the proportion of deaths that are attributable to increased levels of frailty.

**Methods:** We analyzed 42,953 participants from the Screening Across the Lifespan Twin Study (aged 41-95 years at baseline) with up to 20-years’ mortality follow-up. The FI was constructed using 44 health-related items. Deaths due to cardiovascular disease (CVD), respiratory-related causes and cancer were considered in the cause-specific analysis. Generalized survival models were used in the analysis.

**Results:** Increased FI was associated with higher risks of all-cause, CVD, and respiratory-related mortality. No significant associations were observed for cancer mortality. No attenuation of the mortality associations was found in unrelated individuals when adjusting for familial effects in twin pairs. The associations were time-dependent with relatively greater effects observed in younger ages. The proportion of deaths attributable to FI levels >0.10 were 13.0% of all-cause deaths, 14.7% of CVD deaths and 12.5% of respiratory-related deaths in men, and 12.2% of all-cause deaths, 9.9% of CVD deaths and 21.9% of respiratory-related deaths in women.

**Conclusions:** Increased FI predicts higher risks of all-cause, CVD, and respiratory-related mortality independent of familial effects. Increased FI levels have a significant population mortality impact in both men and women.

## Introduction

Frailty is a major health concern associated with aging [1]. It is a state of multisystem physiological decline and inability to maintain homeostasis, gradually leading to an increased risk of multiple adverse outcomes, such as falls, hospitalizations, and death [1]. Frailty is also predictive of poor prognosis and post-operative complications in older surgical patients [2]. The two principal models to operationalize the frailty concept are the Rockwood frailty index (FI) and the Fried phenotypic model (FP) [3]. The FI defines the level of frailty as the ratio of the number of various health deficits present to the number of deficits considered whereas the FP classifies individuals as non-frail, pre-frail or frail based on poor grip strength, slow walking speed, low physical activity, exhaustion, and unintentional weight loss. Being a continuous scale measure, the FI provides good sensitivity even at the lower end of the frailty continuum, which facilitates studies in younger individuals [4, 5].

The FI is associated with all-cause mortality in different populations, independent of other major risk factors [6]. The FI-mortality relationship is well-established, particularly among older individuals (>65 years), but less is known for younger adults and for cause-specific mortality – aspects that may have implications for prevention. Both frailty and risk of death can be influenced by familial background, such as genetic and shared environmental factors, which may confound frailty-mortality associations [7]. However, the extent to which the predictive ability of the FI is affected by familial influences has not been studied. Sex-specific characteristics for the frailty-mortality association have also been suggested, but the matter remains inconclusive [8].

Despite the recognition of frailty as a public health concern [9], its’ direct population mortality impact has received less attention. One way to assess the public health relevance of frailty is to estimate the proportion of deaths that could be avoided if the levels of frailty were decreased in the population. Assessing the fraction of deaths that are attributable to increased frailty, considering both all-cause and cause-specific mortality as outcomes, would provide more precise information about public health relevance.

Consequently, this study aims to estimate the predictive value of the FI on all-cause and cause-specific mortality, separately for men and women. We assess the associations among both unrelated individuals and within twin pairs, to account for familial effects and to assess the degree to which frailty-mortality associations in the general population could be due to shared familial confounding. Secondly, we estimate FI-mortality associations in a time-dependent manner to elucidate the predictive value of FI among the young. Lastly, we investigate the number of deaths that could be attributable to increased population levels of frailty by estimating attributable fractions (AFs) [10–12].

## Methods

### Study population

Data of the present study came from Screening Across the Lifespan Twin Study (SALT), which is part of the Swedish Twin Registry (STR) [13]. In 1998-2002, a total of 44,919 twins born between 1924 and 1958 were interviewed by a structured, computer-assisted telephone interview to collect information about illnesses and health, prescription and nonprescription medication use, occupation, education and lifestyle behaviors. Twins were also asked to go to their local health care centers and provide blood for analyses of clinical chemistries and zygosity determination. Informed consent was obtained from all participants. The study was approved by the Regional Ethics Board in Stockholm (Dnr 2016/1888-31/1).

In the present study, we excluded participants who had more than 20% missing data across the 44 frailty items, and those for whom information on cause-specific mortality could not be retrieved. Eventually, 42,953 twins aged from 41 to 95 years were included in the study, including 11,087 twins whose partner did not respond, 11,548 opposite-sex dizygotic (DZ) twins, 11,812 same-sex DZ twins, and 8,506 monozygotic (MZ) twins. Participants were further grouped into single responders, same-sex dizygotic (DZ) twin pairs, and monozygotic (MZ) twin pairs in the analyses, separately for men and women (Table S1). In the sex-specific analyses, single responders included twins whose partner did not respond in SALT, twins from opposite-sex twin pairs and one randomly selected member of each same-sex twin pair.

### Assessment of the FI

The FI in SALT was constructed based on self-reported data using the Rockwood deficit accumulation model according to standard procedure [4]. Briefly, the deficits to be included in the FI had to have a ≥1% prevalence and ≤10% missingness rate in the study population. Forty-four symptoms, signs, disabilities, and diseases covering a wide range of biological systems and associated with health status were considered in the FI. The items and scoring of the deficits are presented in Table S2. Imputation was used to replace missing values in order to maximize the utilization of the data. Specific methods for the imputation and sensitivity analysis for the imputed data are presented in Appendix S1.

An FI value for each individual was calculated as the number of deficits present divided by the total number of deficits (n=44). For example, an individual having 8 deficits has an FI of 8/44=0.18. To assess the validity of FI in SALT, we examined the distribution and associations with age. The FI was treated as a continuous variable with 10% increase per unit when modeling survival (hazard ratio, HR) and dichotomized to a binary variable by using the cut-off for ‘least fit’ according to Rockwood et al. when estimating the AF: this combines categories ‘relatively fit’ (FI ≤ 0.03) and ‘less fit’ (0.03 < FI ≤ 0.10) as unexposed and categories ‘least fit’ (0.10 < FI ≤ 0.21), ‘frail’ (0.21 < FI ≤ 0.45) and ‘most frail’ (FI ≥ 0.45) as exposed [14]. The selected cut-off for the exposed group for the AF analysis is also close to our cohort median FI of 0.108.

### Assessment of mortality

All-cause mortality data, including dates of deaths, were obtained from linkages of the STR to Swedish national registers through the personal identification number assigned to all residents. All-cause mortality data used in this study were updated on December 31, 2017, yielding up to 20 years of follow-up.

Cause-specific mortality data were obtained from the Cause of Death Register (CDR), with the latest update on December 31, 2014, yielding up to 17 years of follow-up. The CDR records include information about the underlying and contributory causes of death for all individuals registered as Swedish residents in the year of their death. Causes of death were recorded using the International Classification of Diseases (ICD) codes, with ICD-10 from 1997 and onwards. We considered cardiovascular disease (CVD; including stroke), respiratory-related causes, and cancer as the specific causes of death. The ICD codes included in each cause and the consensus classification used when more than one cause of death was recorded are presented in Tables S3 and S4, respectively.

### Assessment of covariates

Years of education, tobacco use status, body-mass index (BMI) and history of diseases were assessed from the self-reported data in SALT. Tobacco status was categorized as non-user (reference category) or user if the person was currently smoking or using smokeless tobacco regularly, or had previously smoked or used smokeless tobacco regularly. BMI was calculated as weight divided by height squared (kg/m2). Individuals were classified as having a history of CVD, chronic respiratory diseases, and/or cancer, if he/she reported any of the following conditions respectively: 1) angina pectoris, myocardial infarction, heart failure, stroke, high blood pressure, claudication, irregular cardiac rhythm/atrial fibrillation, circulation problems in limbs, or thrombosis; 2) asthma, or chronic lung disease; 3) cancer, tumor disease, or leukemia.

### Statistical methods

We used generalized survival models (GSM) for the association between FI and mortality [15, 16]. Briefly, these are a generalization of flexible parametric survival models where the underlying baseline hazard is fitted as a smooth spline term, in our case via a natural cubic spline with 3 to 5 degrees of freedom for different outcomes and cohorts. In all cases, we used attained age as the underlying time scale, and censored subjects in the cause-specific analyses at the date when they died from other causes than the current outcome of interest or the end of follow-up.

We first tested the association assuming a time-constant effect among single responders, same-sex DZ twin pairs, and MZ twin pairs (see Table S1 for grouping). Next, we tested if there were significant time-dependent effects for those causes of death that were significant in the time-constant model among single responders. To elucidate sex-specific characteristics of the associations, all analyses were performed separately for men and women. Years of education, tobacco use, and BMI were included as covariates in all models. For the cause-specific analyses, history of cancer, CVD, or chronic respiratory diseases at baseline was included as a covariate when the corresponding cause of death was the outcome of interest, see Appendix S2, Eq.1, for the basic models for unrelated individuals.

For the same-sex DZ and MZ twin pairs, a between-within decomposition along with a random effect from a gamma distribution were introduced to the GSM, which allow us to adjust for shared familial effects both due to shared patterns of exposure and confounders included in the model as well as general unmeasured similarity in survival patterns in twin pairs [17, 18]. We have previously used a similar model in twin data for telomere length and survival [19]. The model is described in detail in Appendix S2, Eq. 2.

For the subgroup of single responders, we also included interaction terms that allow the association between FI and survival to vary with age at assessment as well as time since assessment; details are presented in Appendix S2, Eq. 3.

The AF was used to quantify the public health impact of FI on the survival outcome in the cohort of single responders based on a time-constant survival model [10–12]. The AF is an integrated predictive measure that takes into account both the prevalence of a risk factor in the population as well as the strength of association between risk factor and outcome. In this study, we report the AFs as the proportion of deaths before attained age of 80 years that could be delayed, or theoretically avoided, if all FI values greater than 0.10 (i.e., the threshold for “least fit” [14]) were reduced to <0.10 in the population.

As our definition of FI includes items related to CVD, respiratory diseases and cancer, we created three “reduced” FIs that were stripped of either CVD-related items (angina pectoris, myocardial infarction, heart failure, stroke, high blood pressure, claudication, irregular cardiac rhythm/atrial fibrillation, circulation problems in limbs, and thrombosis), respiratory-related items (chronic lung diseases and asthma) or cancer. These reduced FIs were then used as exposures in a sensitivity analysis for the corresponding cause-specific survival models. An additional sensitivity analysis using the original 44-item FI was performed for respiratory-related mortality by including only the 17,905 non-tobacco users in the analysis.

P-values reported are two-sided, and the statistical significance level was set at p<0.05. All analyses were conducted using Stata 15.1 and R 3.4.3. Statistical analyses involving GSMs were implemented using the rstpm2 package [20].

## Results

Characteristics of the study population are presented in Table 1. Of 42,953 individuals, 46.4% were men and 47.3% were complete twin pairs. At baseline, the median level of FI was 0.097 in men, and 0.119 in women. FI distributions were skewed with a long right tail in all groups (total cohort, males, and females) (Figure S1). The single responders tended to be older, had a lower education, and used tobacco products more frequently (in men only) than those from complete pairs.

**Table 1.** Characteristics of the study population

|  | Men (N=19,924) |  |  | Women (N=23,029) |  |  |
| --- | --- | --- | --- | --- | --- | --- |
|  | Single responders | DZ twins | MZ twins | Single responders | DZ twins | MZ twins |
| Number of participants | 15,473 | 5,266 | 3,636 | 17,321 | 6,546 | 4,870 |
| Age at baseline, mean (SD) | 58.7 (10.6) | 57.4 (9.6) | 56.9 (9.2) | 59.6 (11.2) | 58.5 (10.3) | 57.8 (10.2) |
| BMI (kg/m<sup>2</sup>), mean (SD) | 25.5 (3.1) | 25.6 (3.1) | 25.5 (3) | 24.6 (3.7) | 24.5 (3.7) | 24.4 (3.8) |
| Education (year), mean (SD) | 10.4 (3.2) | 10.6 (3.2) | 10.8 (3.2) | 10.4 (3.2) | 10.5 (3.2) | 10.8 (3.2) |
| Tobacco products use, % | 64.3 | 63.1 | 59.5 | 53.9 | 54.3 | 52.5 |
| FI, median (IQR) | 0.097<br>(0.102) | 0.094<br>(0.097) | 0.091<br>(0.097) | 0.125<br>(0.125) | 0.119<br>(0.125) | 0.119<br>(0.119) |
| History of diseases at baseline, % |  |  |  |  |  |  |
| CVD | 35.5 | 34.0 | 31.5 | 37.4 | 35.6 | 34.8 |
| Respiratory disease | 9.6 | 8.2 | 9.6 | 11.9 | 10.8 | 11.7 |
| Cancer | 4.7 | 4.2 | 4.7 | 9.1 | 8.5 | 9.3 |
| Time to follow-up, median (IQR) |  |  |  |  |  |  |
| All-cause mortality | 16.7 (2.8) | 16.9 (2.4) | 16.9 (2.3) | 16.9 (2.6) | 17.0 (2.5) | 17.1 (2.4) |
| Cause-specific mortality | 14.0 (2.5) | 14.1 (2.3) | 14.1 (2.3) | 14.2 (2.5) | 14.3 (2.2) | 14.3 (2.3) |
| Death during the follow up, % |  |  |  |  |  |  |
| All causes | 31.3 | 26.0 | 23.4 | 28.3 | 25.0 | 22.1 |
| CVD | 8.9 | 7.0 | 5.7 | 7.4 | 6.1 | 4.8 |
| Respiratory-related causes | 2.7 | 2.1 | 1.9 | 2.4 | 2.2 | 1.7 |
| Cancer | 9.0 | 7.7 | 7.5 | 7.0 | 6.7 | 5.9 |

Note. Single responders included twins whose partner did not respond in SALT, twins from opposite-sex twin pairs and one randomly selected member of each same-sex twin pair; DZ twins refer to same-sex DZ twins.
Participants who used tobacco products include current smokers, ex-smokers, and snuff users at baseline.
Abbreviations: SD: standard deviation; BMI: body mass index; FI: frailty index; IQR: interquartile range; CVD: cardiovascular diseases.

During 20 years of follow-up for all-cause mortality and 17 years of follow-up for cause-specific mortality since 1998-2002, 12,222 (28.5%) deaths were recorded, with 3,270 (7.6%), 1,051 (2.4%), and 3,302 (7.7%) deaths due to CVD, respiratory-related causes, and cancer, respectively. After controlling for education, tobacco use, BMI, the history of corresponding diseases, and familial effects in twins, increased FI significantly predicted higher risks of deaths due to all causes, CVD, and respiratory-related causes in time-constant models in both men and women (Figure 1, Table S5). No significant associations were observed for cancer mortality (Figure 1). The associations of FI did not exhibit marked differences among single responders, same-sex DZ twins, and MZ twins and the associations were also similar in men and women (Figure 1). Additionally, we found time-dependent effects of FI for all-cause, CVD, and respiratory-related mortality in both men and women, with relatively greater HRs at younger ages and effect sizes decreasing with increasing age at FI assessment for all the causes (Figure 2, single responders only), with a 1-2% decrease in strength of association per extra year of age at assessment (Table S6; single responders only).

**Figure 1.**
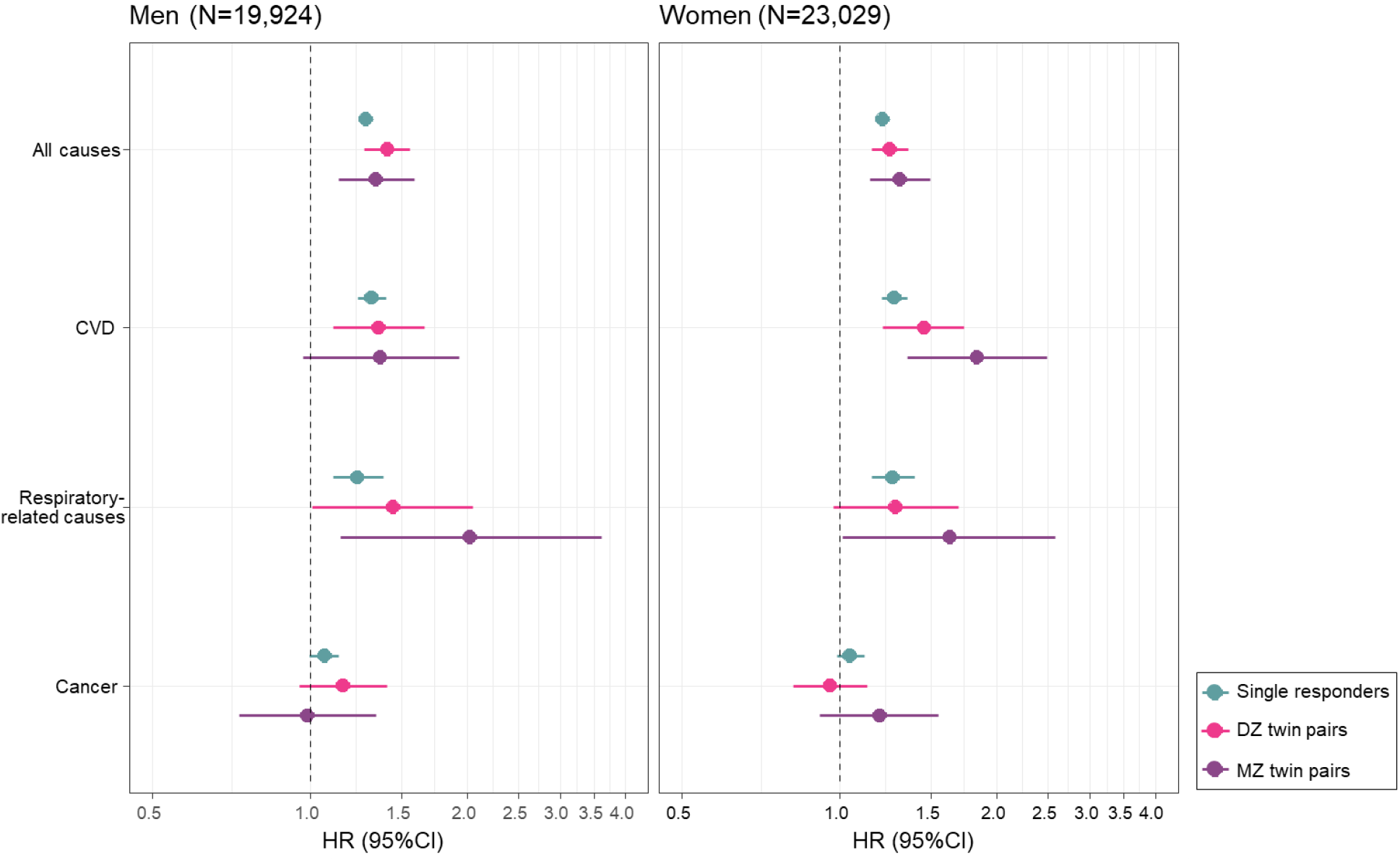
Time-constant effects of 10% increase in the FI on all-cause and cause-specific mortality Increased FI significantly predicted higher risks of deaths due to all causes, CVD, and respiratory-related causes in time-constant models in both men and women. No significant associations were observed for cancer mortality. The associations of FI did not exhibit marked differences among single responders, same-sex DZ twins, and MZ twins and the associations were also similar in men and women. All models considered attained age as time scale, adjusted for BMI, years of education, and tobacco use status; and additionally adjusted for history of CVD, respiratory diseases, or cancer in corresponding cause-specific mortality analysis. Abbreviations: FI: frailty index; CVD: cardiovascular diseases; DZ: dizygotic; MZ: monozygotic.

**Figure 2.**
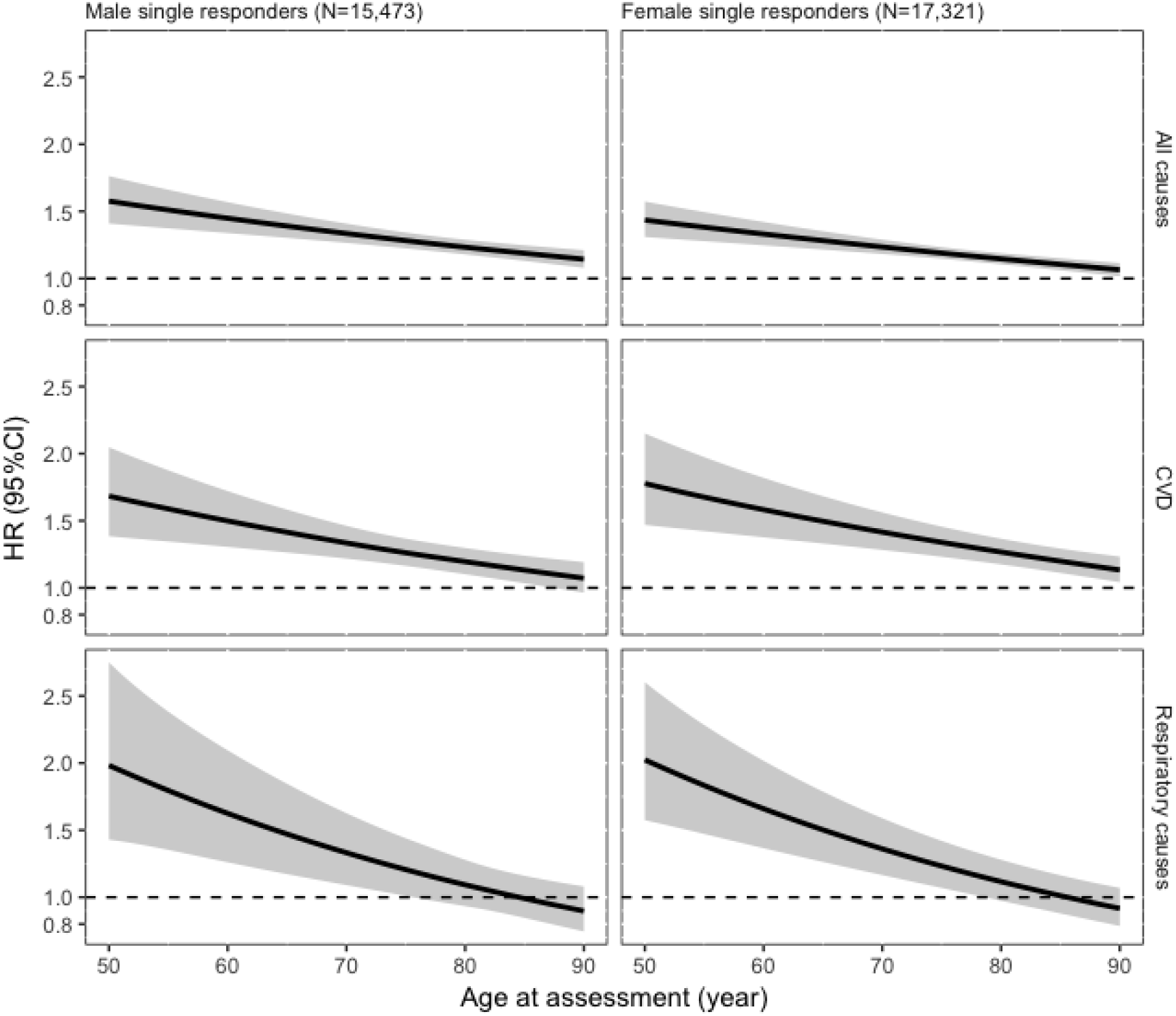
Time-dependent effects of 10% increase in the FI on all-cause and cause-specific mortality in single responders Time-dependent effects of FI for all-cause, CVD, and respiratory-related mortality in both men and women were observed, with relatively greater HRs at younger ages and effect sizes decreasing with increasing age at FI assessment for all the causes. All models adjusted for BMI, years of education, and tobacco use status; and additionally adjusted for history of CVD, or respiratory diseases in corresponding cause-specific mortality analysis, and were fitted as a function of age at FI assessment. Abbreviations: FI: frailty index; CVD: cardiovascular diseases.

Sensitivity analyses demonstrated that the association of FI and cause-specific mortality did not change appreciably when FI was stripped of item(s) related to the cause of death under examination, or when tobacco users were excluded from the study population (Table S7).

AFs before the age of 80 years are provided in Table 2, and the AF estimates across the whole age continuum are presented in Figure S2. In men, the AFs (95% CI) revealed that 13.0% (10.6% - 15.3%) of all-cause deaths, 14.7% (8.4% - 21.0%) of CVD deaths, and 12.5% (1.6% - 23.3%) of respiratory-related deaths were attributed to FI levels >0.10. In women, the AFs were 12.2% (9.0% - 15.4%) for all-cause deaths, 9.9% (1.5% - 18.4%) for CVD deaths, and 21.9% (7.7% - 36.2%) for respiratory-related deaths. The AFs demonstrated a decreasing trend toward the oldest ages but were relatively constant before the age of 80 (Figure S2).

**Table 2.** Attributable fractions (AF) for all-cause and cause-specific mortality in single responders before the age of 80.

| Outcomes | Male single responders<br>(N=15,473) | Female single responders<br>(N=17,321) |
| --- | --- | --- |
| All causes | 13.0% (10.6% - 15.3%) | 12.2% (9.0% - 15.4%) |
| CVD | 14.7% (8.4% - 21.0%) | 9.9% (1.5% - 18.4%) |
| Respiratory-related<br>causes | 12.5% (1.6% - 23.3%) | 21.9% (7.7% - 36.2%) |

The AF represents the proportion of deaths that could be avoided if FI levels higher than 0.10 were eliminated from the population.

## Discussion

In this study, we constructed the FI based on the Rockwood cumulative deficit model in a Swedish twin population and assessed its predictive value on all-cause and cause-specific mortality. During the 20-year follow-up, an increase in FI conveyed a significant risk for all-cause, CVD, and respiratory-related mortality, in both men and women. However, no associations with cancer mortality were found. The associations were persistent when adjusted for tobacco use, education and BMI. The time-dependent analysis demonstrated that for all the causes of death, the effects varied with age at FI assessment, with relatively greater HRs observed at younger ages. When adjusting for familial effects in twin pairs, we found no attenuation of the associations between FI and mortality in comparison to those observed in single responders. In fact, even an increasing trend in the strength for FI and respiratory-related mortality in men and for FI and CVD-mortality in women was found when the familial effects were accounted for. The estimated AFs demonstrated that (theoretical) reduction of the FI levels below 0.10 would prevent 11.2-16.5% of all-cause, CVD, and respiratory-related deaths in men and women before the age of 80 years.

Although a wealth of evidence exists to demonstrate that a higher level of frailty is predictive of all-cause mortality, previous studies have mostly been conducted in unrelated individuals [6, 21]. The associations may thus be confounded by familial factors, such as genetics and early life environment, affecting the risk of both frailty and mortality. We tested this by modeling the effects in MZ and DZ twin pairs, and found no evidence that the associations were attenuated compared to the results obtained in single responders. In other words, the FI is at least as useful a predictor of mortality after adjustment for shared family history in twins as in a general, unrelated population, suggesting that FI captures the individual association between frailty at time of assessment and risk of mortality with little or no confounding from shared familial lifestyle or genetic factors.

As frailty is considered to be a manifestation of aging, FI in relation to mortality has been mostly investigated in older populations [6, 21]. Several studies with wide age spectra have nevertheless suggested that a negative effect of increased FI on mortality could also be observed in younger adults [14, 22–24]. In fact, previous findings by others [22] and us [24, 25] have suggested that the frailty-mortality association is relatively stronger among the young than the old. In the present study, we modeled the effect in a time-dependent manner and found that for all causes, higher FI was associated with a greater relative risk of mortality at younger ages and the associations attenuated towards the oldest ages. Such accumulating evidence suggests that a shift towards more research into frailty in younger populations is warranted.

Frailty presents a sex-specific pattern that is similar to the male-female health-survival paradox in which women experience higher rates of disability and comorbidity yet still outlive men [26]. Women have higher levels of frailty throughout the age range, but men are more vulnerable to death at any given level of frailty [4, 8]. However, whether frailty is a stronger risk factor for men than for women is currently unclear and different studies have reported contrasting results [27–31]. We tested this hypothesis by assessing the associations with both all-cause and cause-specific mortality in men and women separately. Overall, the HRs were similar in men and women, even when adjusted for familial effects in twins. The time-dependent effects also demonstrated similar patterns in men and women, with relatively greater HRs observed at midlife and younger old ages and an attenuating trend towards the oldest ages.

Other studies that used the Fried phenotypic model instead of FI to measure frailty have also demonstrated that increased frailty is predictive of CVD mortality: Crow et al reported an association that was adjusted for sex in an analysis including men and women [32], and Veronese et al. analyzed men and women separately and demonstrated that the association was stronger in women [33]. However, as analyses into cause-specific mortality and frailty are currently limited, more research is warranted to establish the relationship between frailty and cause-specific mortality.

Due to its association with various adverse outcomes, frailty presents a public health concern. The risk is not restricted to the highest end of the frailty continuum; lower levels of frailty, such as the pre-frailty state measured using the FP, are also predictive of mortality [3]. Therefore, we assessed the proportion of deaths that could be avoided by decreasing the levels of frailty using the cut-off for “least fit” according to Rockwood et al [14]. We found that before the age of 80, these proportions are slightly greater than 10% for all the causes investigated, in both men and women. Our AF estimates for all-cause mortality are higher than previously reported in a systematic review that estimated the population risk of all-cause mortality attributable to frailty to be 3-5%. The systematic review used a higher cut-off, corresponding to the frailty state, and the individuals included were older than 65 years [21]. These differences may account at least in part for the differing results. In fact, the AFs in our study showed a decreasing trend toward the very old ages, indicating that higher frailty bears a relatively greater population risk of mortality at midlife and younger old ages. A recommendation to screen all individuals at 70 years and older at health care facilities has been put forward [34]. However, pertaining to mortality risk stratification, our results would suggest a potential benefit of considering also middle-aged individuals for frailty screening.

The present study has several strengths. First, we had a large population of twins that allowed us to control for unmeasured familial effects. Second, contrary to previous research mainly focusing on FI among older individuals (>65 years), our cohort also included younger adults and had an age range from 40 to 95 years, which allowed us to model the association between frailty and mortality as a continuous function of age at assessment. Third, we had a very long follow-up, up to 20 years for all-cause mortality and up to 17 years for cause-specific mortality, which enables us to draw conclusions on the long-term predictive ability of the FI. Indeed, the associations in this study demonstrate that the FI is predictive of mortality in a long-term follow-up and align with a previous study by our group, in which we demonstrated that the FI is predictive of all-cause mortality for up to 30 years [24].

The present study also has some limitations. Frailty was measured using only one scale, the FI, and was based on self-reported data. Our FI contained items of cancer, CVD, and chronic respiratory diseases. Hence, in the cause-specific analysis, we adjusted each of the analyses for the history of the given disease that was the cause of death in that analysis. For each of the cause-specific mortality analyses, we additionally created a modified FI that was stripped of item(s) related to the cause of death under examination. Lastly, we repeated the analysis of respiratory-related mortality in non-tobacco users. The results remained unchanged in these sensitivity analyses, suggesting that the FI is a robust marker for predicting risk of death due to respiratory-related causes and CVD, even in the presence of these diseases.

In conclusion, an increase in FI predicted higher risks of all-cause, CVD, and respiratory-related mortality, independent of familial effects, and exhibited time-dependent associations suggesting that a higher level of frailty is a relatively greater risk factor for middle-aged individuals than for older. Our results also highlight the significant population mortality impact of increased FI levels, including the moderately elevated FI levels that are lower than the “frailty state”.

## Supporting information

## Acknowledgements

Not applicable.

