## Supplementary material for "The frailty index is a predictor of cause-specific mortality independent of familial effects from midlife onwards"

Table S1 Grouping of the study population

| | | 11,087 twins whose partner did not respond | | 11,548 opposite-sex DZ twins | | 11,812 same-sex DZ twins | | 8,506 MZ twins | |
|---|---|---|---|---|---|---|---|---|---|
| | | 5,248 men 0 pairs | 5,839 women 0 pairs | 5,774 men 0 pairs | 5,774 women 0 pairs | 5,266 men 2,633 pairs | 6,546 women 3,273 pairs | 3,636 men 1,818 pairs | 4,870 women 2,435 pairs |
| Grouped in the analyses as: | | | | | | | | | |
| Single responders | Men | √ | | √ | | 50% √ | | 50% √ | |
| | Women | | √ | | √ | | 50% √ | | 50% √ |
| (Same-sex) DZ twin pairs | Men | | | | | √ | | | |
| | Women | | | | | | √ | | |
| MZ twin pairs | Men | | | | | | | √ | |
| | Women | | | | | | | | √ |

Single responders included twins whose partner did not respond in SALT, twins from opposite-sex twin pairs and one randomly selected member of each same-sex twin pair.
DZ: dizygotic twin; MZ: monozygotic twin

Table S2 44 frailty items and the coding rules

| No. | Questions | Coding |
|---|---|---|
| 1 | How do you estimate your general health? | Excellent=0, Good=0.25, Average=0.5, Not so good=0.75, Bad=1 |
| 2 | Do you think your health status prevents you from doing things you want to do? | Not at all=0, To some extent=0.5, A great deal=0 |
| 3 | How many times a year do you get serious infections (other than respiratory)? | 0-1 times=0, 2-4 times=0.5, 5 times or more =1 |
| 4 | Do you have buzzing in the ears? | Both ears or one ear=1, No=0 |
| 5 | Do you have or have you had angina pectoris | No=0, Yes=1 |
| 6 | Do you have or have you had heart attack | No=0, Yes=1 |
| 7 | Do you have or have you had heart failure | No=0, Yes=1 |
| 8 | Do you have or have you had high blood pressure | No=0, Yes=1 |
| 9 | Do you have or have you had lipid disorder, for example high cholesterol or high triglycerides | No=0, Yes=1 |
| 10 | Do you have or have you had vascular spasm in the legs (intermittent claudication) | No=0, Yes=1 |
| 11 | Do you have or have you had clot in the leg (venous thrombosis) | No=0, Yes=1 |
| 12 | Do you have or have you had cerebral hemorrhage or clot in the brain (stroke) | No=0, Yes=1 |
| 13 | Do you have or have you had TIA attacks (temporary weakness or paralysis or reduction of sensibility) | No=0, Yes=1 |
| 14 | Do you have or have you had irregular cardiac rhythm/atrial fibrillation | No=0, Yes=1 |
| 15 | Do you have or have you had chronic lung disease (including chronic bronchitis and emphysema) | No=0, Yes=1 |
| 16 | Do you have or have you had dizziness | No=0, Yes=1 |
| 17 | Do you have or have you had rheumatoid arthritis | No=0, Yes=1 |
| 18 | Do you have or have you had knee joint problem | No=0, Yes=1 |
| 19 | Do you have or have you had sciatica | No=0, Yes=1 |
| 20 | Do you have or have you had osteoporosis | No=0, Yes=1 |
| 21 | Do you have or have you had hip joint problem | No=0, Yes=1 |

| | | |
|---|---|---|
| 22 | Do you have or have you had back pain | No=0, Yes=1 |
| 23 | Do you have or have you had neck pain | No=0, Yes=1 |
| 24 | Do you have or have you had diabetes (including old age diabetes, and excluding pregnancy diabetes) | No=0, Yes=1 |
| 25 | Do you have or have you had goiter | No=0, Yes=1 |
| 26 | Do you have or have you had glandular diseases (excluding goiter) | No=0, Yes=1 |
| 27 | Do you have or have you had gall bladder problem | No=0, Yes=1 |
| 28 | Do you have or have you had liver disease (for example, cirrhosis) | No=0, Yes=1 |
| 29 | Do you have or have you had gout | No=0, Yes=1 |
| 30 | Do you have or have you had kidney disease | No=0, Yes=1 |
| 31 | Do you have or have you had stomach or intestine problems | No=0, Yes=1 |
| 32 | Do you have or have you had recurring urinary tract problems | No=0, Yes=1 |
| 33 | Do you have or have you had cancer, tumor disease or leukemia | No=0, Yes=1 |
| 34 | Do you have or have you had migraine | No=0, Yes=1 |
| 35 | Do you have or have you had asthma | No=0, Yes=1 |
| 36 | Do you have or have you had allergy | No=0, Yes=1 |
| 37 | Do you have recurrent periods of coughing? | No=0, Yes=1 |
| 38 | You felt depressed. Never, seldom, often or always during the past week? | Never or almost never=0, Seldom=0.5, Often, always or almost always=1 |
| 39 | You were happy. Never, seldom, often or always during the past week? | Never or almost never=1, Seldom=0.5, Often, always or almost always=0 |
| 40 | You felt lonely. Never, seldom, often or always during the past week? | Never or almost never=0, Seldom=0.5, Often, always or almost always=1 |
| 41 | Do you have or have you had any physical handicap | No=0, Yes=1 |
| 42 | Do you have or have you had Crohn's disease or Ulcerative colitis | No=0, Yes=1 |
| 43 | How is your vision? | Good=0, Reduced=0.5, Highly reduced or blind=1 |
| 44 | How is your hearing? | Good=0, Reduced=0.5, Highly reduced=1 |

Table S3 ICD codes used to classify the cause-specific mortality.

| | **ICD-10** |
|---|---|
| **Non-stroke CVD** | I20-I25 |
| | I70 |
| | I73.9 |
| **Stroke** | I60-I61 |
| | I63-I64 |
| **Cancer** | C00-C97 |
| | B21 |
| **Respiratory-related causes** | J00-J99 |

Non-stoke CVD and Stroke were considered as CVD-mortality.
ICD: International Classification of Diseases; CVD: cardiovascular disease.

Table S4 Consensus classification for the cause-specific mortality when more than one cause of death was recorded.

| **Cancer** | **CVD** | **Respiratory-related causes** | **Consensus** | **Count in men** | **Count in women** |
|---|---|---|---|---|---|
| - | - | + | Respiratory-related causes | 586 | 571 |
| - | + | - | CVD | 1,075 | 983 |
| + | - | - | Cancer | 1,278 | 1,339 |
| - | + | + | CVD | 332 | 273 |
| + | + | - | Cancer | 168 | 74 |
| + | - | + | Cancer | 222 | 142 |
| + | + | + | Cancer | 57 | 22 |

CVD: cardiovascular disease.

Table S5 Time-constant Effects of 10% FI increase on all-cause and cause-specific mortality.

| | Men (N=19,924) | | | Women (N=23,029) | | |
|---|---|---|---|---|---|---|
| | Single responders | DZ twins | MZ twins | Single responders | DZ twins | MZ twins |
| All causes | 1.28 (1.24, 1.32) | 1.40 (1.27, 1.55) | 1.34 (1.13, 1.58) | 1.21 (1.18, 1.25) | 1.25 (1.15, 1.35) | 1.30 (1.14, 1.49) |
| CVD | 1.31 (1.23, 1.40) | 1.35 (1.11, 1.66) | 1.37 (0.97, 1.92) | 1.27 (1.15, 1.34) | 1.45 (1.21, 1.73) | 1.83 (1.35, 2.49) |
| Respiratory-related causes | 1.23 (1.11, 1.38) | 1.44 (1.01, 2.05) | 2.03 (1.14, 3.60) | 1.26 (1.15, 1.39) | 1.28 (0.97, 1.69) | 1.62 (1.02, 2.58) |
| Cancer | 1.06 (1.00, 1.14) | 1.15 (0.95, 1.40) | 0.99 (0.73, 1.34) | 1.05 (0.99, 1.11) | 0.96 (0.81, 1.13) | 1.19 (0.92, 1.55) |

All models considered attained age as time scale, adjusted for BMI, years of education, and tobacco use status, and additionally adjusted for history of CVD, respiratory diseases, or cancer in corresponding cause-specific mortality analysis.

FI: frailty index; CVD: cardiovascular diseases; DZ: dizygotic; MZ: monozygotic.

Table S6 Assessment of time-dependent associations between FI and mortality in single responders: mortality hazard ratios for an increase in age at FI assessment by one year; i.e. $\exp(\beta_{age})$ in Appendix S2, Eq. 3

| | Male single responders (N=15,473) | | Female single responders (N=17,321) | |
|---|---|---|---|---|
| | HR (95%CI) | P value (Wald) | HR (95%CI) | P value (Wald) |
| All causes | 0.99 (0.99, 1.00) | 0.002 | 0.99 (0.99, 0.99) | <0.001 |
| CVD | 0.99 (0.98, 0.99) | <0.001 | 0.99 (0.98, 0.99) | <0.002 |
| Respiratory-related causes | 0.98 (0.97, 0.99) | 0.002 | 0.98 (0.97, 0.99) | <0.003 |

Age at FI assessment was included as continuous variable in the models

Table S7 Sensitivity analyses of effect of 10% FI increase on all-cause and cause-specific mortality in single responders

| | | Male single responders (N=15,473) | Female single responders (N=17,321) |
|---|---|---|---|
| CVD | Analysis a | 1.18 (1.11, 1.25) | 1.17 (1.11, 1.23) |
| Respiratory-related causes | Analysis b | 1.21 (1.09, 1.35) | 1.24 (1.13, 1.35) |
| | Analysis c | 1.28 (1.04,1.58) | 1.23 (1.07, 1.42) |
| Cancer | Analysis d | 1.06 (1.00, 1.13) | 1.05 (0.99, 1.11) |

Analysis a: Removed 9 CVD-related conditions from 44 frailty items, including angina pectoris, myocardial infarction, heart failure, stroke, high blood pressure, claudication, irregular cardiac rhythm/atrial fibrillation, circulation problems in limbs, and thrombosis.
Analysis b: Removed 2 respiratory-related conditions from 44 frailty items, including chronic lung diseases, and asthma.
Analysis c: Excluded 25,048 tobacco users from the present study, leaving 17,905 participants.
CVD: cardiovascular diseases.
Analysis d: Removed cancer from 44 frailty items.
All models considered attained age as time scale, adjusted for BMI, years of education, tobacco use status, and history of CVD, respiratory diseases, or cancer in corresponding cause-specific mortality analysis assuming a time-constant effect.

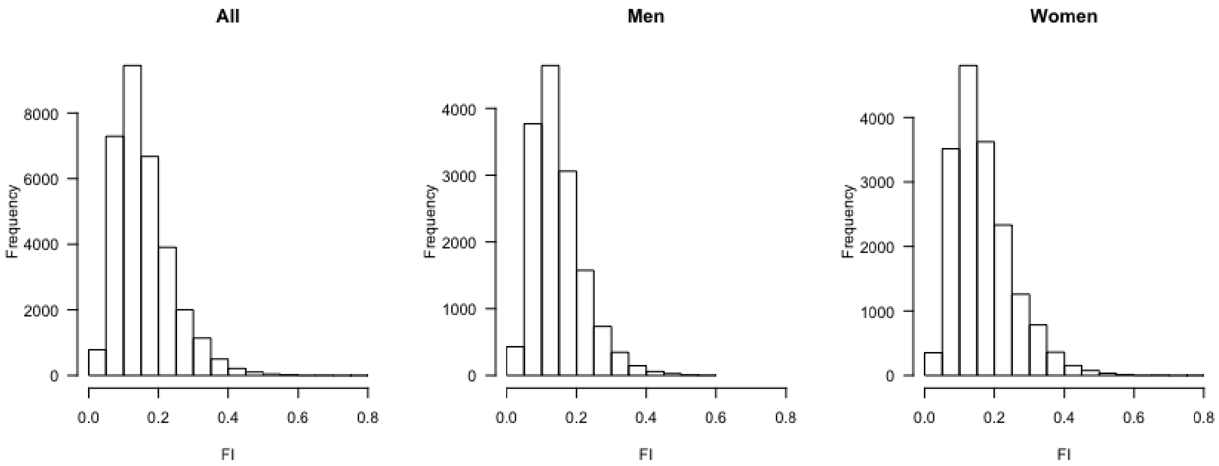

Figure S1 Distribution of the frailty index in our sample.

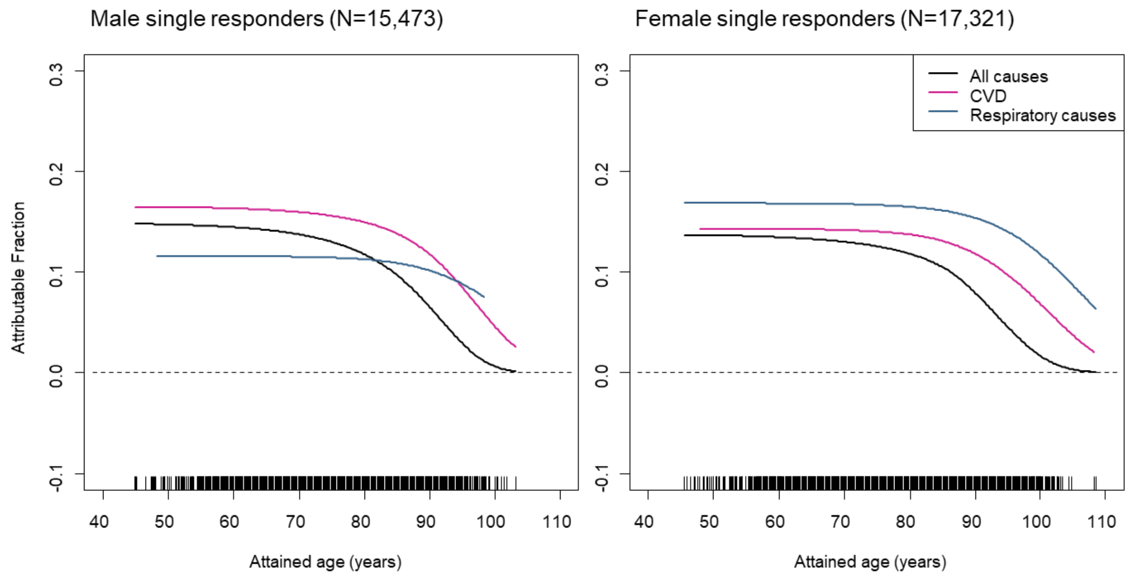


Figure S2 Attributable fraction (AF) functions for all-cause and cause-specific mortality in single responders.

The AF represents the proportion of deaths that could be avoided if FI levels higher than 0.10 were eliminated from the population as a function of attained age.

### Appendix S1 Imputation of missing values

After selection of the 44 items, we imputed missing values by chained equations, using logistic regression for binary variables and predictive mean matching for categorical variables and continuous variables. When a certain frailty item was imputed, the other 43 frailty items, age, sex, BMI, education level, and smoking status were used as predictors in the regression models. Ten rounds of imputations were performed and the pooled mean from the simulations was used as the final value for each missing data point. Next, all the item scores were summed and the FI was calculated by dividing the sum by 44.

We performed sensitivity analyses of FI and all-cause mortality for those individuals with no missing data in the FI items (n=40,171) and similar results were observed with those from the imputed dataset (n=42,953). (Data not shown)

Appendix S2 Specification of generalized survival models (GSM) fitted to the data

The basic GSM for all-cause mortality in single responders was specified with a log-log link function and attained age as the underlying time scale. Consequently, the model can be written as

$$\log\left(-\log\left(S(t_i|x_i)\right)\right) = s_0(t_i;\gamma) + \beta_1 FI_i + \beta_2 Edu_i + \beta_3 b_i + \beta_4 BMI_i \qquad (1)$$

where $S(t_i|x_i)$ is the conditional survival function evaluated at time $t_i$ and the covariate pattern $x_i$ observed for the $i$-th individual in the data set, and $s_0(t_i;\gamma)$ is a natural cubic spline with $\gamma$ degrees of freedom, representing the baseline hazard of mortality as a function of attained age. $FI_i$, $Edu_i$, $b_i$ and $BMI_i$ represent the covariates FI (in 10% increments), education in years, binary tobacco use and BMI in kg/m2, respectively, as observed for the $i$-th individual. For cause-specific outcomes, this model contains an extra term $\beta_5 Hist_i$, which is a binary indicator for a previous history of the disease that is the cause of interest. We report the respective estimated $\exp(\beta_1)$as hazard ratios for FI in single responders in Figure 1 and Table S6.

For both DZ and MZ twins, this model was reformulated as a between-within model with a shared gamma random effect (what is commonly in survival analysis referred to as "gamma frailty", an expression that we avoid for the obvious potential for confusion here). In this model, the effects of covariates are split into a between-pair effect $\beta_B$ that captures covariate effects shared by both members of the twin pair, and a within-pair effect $\beta_W$ that captures covariate effects specific to individuals and adjusted for shared effects. In addition, the model includes a random gamma effect that captures shared similarity in survival between twins in a pair that is independent of the included covariates, see Ref 17 in the main paper. Similar to Ref 18 in the main paper, the resulting extended GSM for all-cause survival can be written as

$$\log\left(-\log\left(S\left(t_{ij}|x_{ij},u_i\right)\right)\right) = s_0\left(t_{ij};\gamma\right) + \log(u_i) + \beta_B\overline{FI}_i + \beta_W FI_{ij} + \delta_{B_m}\bar{C}_{i_m} + \delta_{W_m}C_{ij_m} \qquad (2)$$

Here $S\left(t_{ij}|x_{ij},u_i\right)$ is again the survival function, but this time evaluated at time $t_{ij}$ and covariate pattern $x_{ij}$ observed for the $j$-th twin ($j \in 1:2$) in the $i$-th twin pair ($i$ ranging from 1 to the number of twin pairs in the analysis), and conditioned on the random effect $u_i$ drawn from a gamma distribution and shared between the member of the $i$-th twin pair. As before, $s_0\left(t_{ij};\gamma\right)$ is a natural cubic spline term capturing the baseline hazard. The main exposure FI is included via the between-pair effect $\beta_B$, scaling the contribution of the averaged value $\overline{FI}_i$ in the $i$-th twin pair, and the within-pair effect $\beta_W$, scaling the observed value $FI_{ij}$ for twin $j$ in pair $i$. Adjustment for the other covariates follows the same split and is indicated in the equation via the averaged between-pair term $\bar{C}_{i_m}$ and the observed individual term $C_{ij_m}$, where $m$ varies over education, tobacco use, BMI and (for cause-specific analyses) disease history, as for the singleton model above. We report the respective estimated $\exp(\beta_W)$ in Figure 1 and Table S6 as hazard ratios for FI in DZ and MZ twins, adjusted for shared familial effects.

In order to allow the effect of FI to vary with age at measurement in a consistent manner, we split a simple time-varying effect of FI expressed as an interaction with the underlying time scale into two components:

$$t_i FI_i = a_i FI_i + (t_i - a_i) FI_i$$

where as above, $t_i$ is the attained age at the end of followup and $FI_i$ the value of FI observed at the start of follow-up for the $i$-th subject in the data set; the new term $a_i$ is the age of the subject at measurement of FI, which is just the age at start of follow-up. Consequently, this split allows to add separate effects for age at measurement and time since measurement with regard to the association between FI and survival. The corresponding full model can be written as

$$\log\left(-\log\left(S(t_i|x_i)\right)\right) = s_0(t_i;\gamma) + \beta_{fix} FI_i + \beta_{age}(a_i - 70) FI_i + \beta_{ts}(t_i - a_i) FI_i + \beta_2 Edu_i + \beta_3 Tob_i + \beta_4 BMI_i \quad (3)$$

Here, $\beta_{ts}$ is an interaction term that models how the association between FI and survival varies with time since measurement, $\beta_{age}$ is the same for age at measurement, and $\beta_{fix}$ is the main effect capturing the association between FI and survival when measured at age 70 and at time of measurement (which is when the interaction terms are zero). We report the respective estimated $\exp(\beta_{age})$ as hazard ratios for FI in Table S2, and plot the product with the estimated $\exp(\beta_{fix})$ in Figure 2 for the time-varying hazard ratio as function of age at measurement.
